# The community and ecosystem consequences of intraspecific diversity: a meta-analysis

**DOI:** 10.1101/328112

**Authors:** Allan Raffard, Frédéric Santoul, Julien Cucherousset, Simon Blanchet

## Abstract

Understanding the relationships between biodiversity and ecosystem functioning has major implications. Biodiversity–ecosystem functioning relationships are generally investigated at the interspecific level, although intraspecific diversity (i.e. within-species diversity) is increasingly perceived as an important ecological facet of biodiversity. Here, we provide a quantitative and integrative synthesis testing, across diverse plant and animal species, whether intraspecific diversity is a major driver of community dynamics and ecosystem functioning. We specifically tested (*i*) whether the number of genotypes/phenotypes (i.e. intraspecific *richness*) or the specific identity of genotypes/phenotypes (i.e. intraspecific *variation*) in populations modulate the structure of communities and the functioning of ecosystems, (*ii*) whether the ecological effects of intraspecific richness and variation are strong in magnitude, and (*iii*) whether these effects vary among taxonomic groups and ecological responses. We found a non-linear relationship between intraspecific richness and community and ecosystem dynamics that follows a saturating curve shape, as observed for biodiversity–function relationships measured at the interspecific level. Importantly, intraspecific richness modulated ecological dynamics with a magnitude that was equal to that previously reported for interspecific richness. Our results further confirm, based on a database containing more than 50 species, that intraspecific variation also has substantial effects on ecological dynamics. We demonstrated that the effects of intraspecific variation are twice as high as expected by chance, and that they might have been underestimated previously. Finally, we found that the ecological effects of intraspecific variation are not homogeneous and are actually stronger when intraspecific variation is manipulated in primary producers than in consumer species, and when they are measured at the ecosystem rather than at the community level. Overall, we demonstrated that the two facets of intraspecific diversity (richness and variation) can both strongly affect community and ecosystem dynamics, which reveals the pivotal role of within-species biodiversity for understanding ecological dynamics.

## I. INTRODUCTION

Understanding the relationships between biodiversity and ecosystem functioning is an intensely active field of research informing on the services provided by biodiversity (Chapin *et al.*, 2000; Loreau, 2000; Hooper *et al.*, 2005; Cardinale *et al.*, 2012). Biodiversity is generally quantified as the taxonomic, functional and/or phylogenetic diversity of a species assemblage, and most studies on biodiversity–ecosystem functioning relationships have to date focused on the interspecific facet of biodiversity (Naeem *et al.*, 1994; Downing & Leibold, 2002; Hillebrand & Matthiessen, 2009). However, biodiversity also includes an intraspecific facet that is defined as the phenotypic, functional and genetic diversity measured within a single species (Odling-Smee, Laland & Feldman, 2003; Bolnick *et al.*, 2003). During the last two decades, intraspecific diversity has been demonstrated to account for a non-negligible part of the total biodiversity measured in plants and animals (Fridley & Grime, 2010; de Bello *et al.*, 2011), representing in some cases up to a quarter of the total variability measured in communities (Fridley & Grime, 2010; de Bello *et al.*, 2011; Siefert *et al.*, 2015).

In parallel, the hypothesis that intraspecific diversity may affect ecological dynamics at levels higher than the population level (for instance the composition and the dynamics of communities and/or the dynamics of ecosystem functions) has been conceptualized (Bolnick *et al.*, 2003, 2011; Hughes *et al.*, 2008; Bailey *et al.*, 2009; Violle *et al.*, 2012). These conceptual insights have been validated by several key experiments both in plants and animals (Whitham *et al.*, 2003; Madritch, Greene & Lindroth, 2009; Matthews *et al.*, 2016; Rudman & Schluter, 2016). For instance, the experimental manipulation of fish phenotypes from several evolutionarily independent fish populations has been shown to generate significant changes in both the community structure of invertebrate prey and the primary productivity of the ecosystem (Harmon *et al.*, 2009; Matthews *et al.*, 2016).

Intraspecific diversity can be characterized based on the *richness* of populations, which corresponds to the differences in the number of genotypes and/or phenotypes composing populations. For instance, populations are often characterized according to their ‘allelic, genotypic or phenotypic richness’, which is a population parallel of species richness, a common metric measured at the interspecific level and classically used to investigate biodiversity–ecosystem function (BEF) relationships (Crutsinger *et al.*, 2006). Intraspecific richness can also affect ecological dynamics hence generating ‘intraspecific BEF’ (Whitham *et al.*, 2006; Crutsinger *et al.*, 2006). The basic hypothesis for intraspecific BEF is that increasing the number of genotypes/phenotypes in a population should alter (either negatively or positively) key ecological functions such as the decomposition rate of organic matter or the structure of communities. For instance, experiments manipulating the number of genotypes (from one to 12 genotypes) in plant (*Solidago altissima*) populations have shown that richer populations contained a higher diversity of invertebrates (Crutsinger *et al.*, 2006). Actually, the ecological consequences of intraspecific richness should follow a saturating curve (i.e. a rapid increase followed by a plateau) as often described for BEF observed at the interspecific level (Hughes *et al.*, 2008). Although rarely tested empirically, this saturating shape could be due to the combined effects of several mechanisms. Populations with different richness could have different ecological consequences because of ecological complementarity among genotypes/phenotypes (i.e. niche partitioning, facilitation occurring when genotypes use different resources), inhibition among genotypes/phenotypes (when multiple genotypes are in competition for resources), or functional redundancy among genotypes/phenotypes that can make populations ecologically equivalent (Johnson, Lajeunesse & Agrawal, 2006; Hughes *et al.*, 2008). Yet, the shape of the relationship between intraspecific richness and ecological dynamics has rarely been investigated empirically and to our knowledge has never been quantified across species.

The ecological consequences of intraspecific diversity can also be investigated through the lens of *variation* in genotypic or phenotypic attributes. Adaptive and non-adaptive evolutionary processes such as natural selection, plasticity or genetic drift can generate unique phenotypic differences among populations. These differences can be associated to key functional processes such as food acquisition or nutrient cycling (e.g. Grant & Grant, 2006; Rudgers & Whitney, 2006; Lowe, Kovach & Allendorf, 2017), resulting in both trophic and non-trophic effects of intraspecific variation on ecological dynamics (Odling-Smee *et al.*, 2003; Whitham *et al.*, 2003; Matthews *et al.*, 2011). For instance, it has been shown experimentally that plant genotypes differing in their susceptibility to herbivores harbour different communities of herbivores (Crutsinger, Cadotte & Sanders, 2009*a*; Barbour *et al.*, 2009*b*). Similarly, mesocosm experiments have shown that differences in diet within predator populations can modify prey community structure (Post *et al.*, 2008; Harmon *et al.*, 2009; Howeth *et al.*, 2013). Non-trophic interactions can also have an important role. For instance, differences in the chemical composition of individuals can result in differences in excretion rate or in leaf chemistry that can then affect ecosystem functions such as primary production or nutrient recycling (Lecerf & Chauvet, 2008; El-Sabaawi *et al.*, 2015). Recently, Des Roches *et al.* (2018) demonstrated that intraspecific variation can affect ecological dynamics to the same extent as the removal or replacement of a species in the environment. Although based on a relatively limited number of studies (25 studies on 15 species), their study confirmed the hypothesis that intraspecific variation might be a non-negligible driver of ecological dynamics.

Here, we investigated – across various species and ecosystems – the extent to which both intraspecific richness and intraspecific variation affect the structure of communities and the functioning of ecosystems, and whether intraspecific diversity is a major driver of ecological dynamics. We reviewed published studies testing the causal effects of intraspecific diversity on ecological dynamics in two meta-analyses synthesizing published data across taxa and ecosystems for intraspecific richness and variation, respectively, and to fulfil three specific objectives. First, we tested the significance and the shape of the relationship between intraspecific richness and ecological dynamics. We expected to find a significant saturating relationship between intraspecific richness and ecological dynamics, because of potential facilitation and functional redundancy among genotypes and phenotypes (Hughes *et al.*, 2008). Second, we tested whether manipulating intraspecific richness has similar effects on ecological dynamics to manipulating interspecific richness, by comparing the ecological effects of intraspecific richness with those of interspecific richness obtained from experimental studies manipulating species richness (Duffy, Godwin & Cardinale, 2017). Finally, we provided a novel and extensive quantitative synthesis testing for the effects of intraspecific variation on ecological dynamics. Des Roches *et al.* (2018) previously focused on studies removing or replacing the target species (by which intraspecific variation was manipulated) to investigate the ecological consequences of intraspecific variation. This strongly restricted the number of available studies for which effects sizes could be calculated, and potentially upwardly biased the resulting estimates (Des Roches *et al.*, 2018). We here relax this restriction by considering all studies manipulating intraspecific variation, and use a null-model approach to provide a more accurate relative effect size of intraspecific variation on ecological dynamics. We also built on this extended data set to partition variance in the ecological consequences of intraspecific variation according to the type of organism manipulated and the type of response variable measured. We tested whether the magnitude of the effects of intraspecific variation on ecological dynamics vary among organism types (primary producers *versus* consumers) and levels of biological organization (community *versus* ecosystem levels). Because primary producers form the basis of trophic chains, we expect stronger ecological effects of intraspecific variation in producers than in consumers. We also expect stronger effects of intraspecific variation on ecosystem functions than on metrics describing community structure because ecosystem functions are affected by both trophic and non-trophic effects of biodiversity (Matthews *et al.*, 2014).

## II. MATERIALS AND METHODS

### (1) Data collection

We compiled data from published articles quantifying the effects of intraspecific diversity in a single species on community structure and/or ecosystem functioning. We focused only on intraspecific diversity that represented the integrative phenotypic effect of multiple evolutionary processes including selection, drift and/or plasticity. As a result, we did not consider articles focusing on experimentally induced intraspecific diversity through induced plastic responses to particular predatory or environmental cues [for example see Werner & Peacor (2003) for a review]. We reviewed several experimental studies manipulating intraspecific variation and/or richness within a single species to test their respective ecological effects. We also reviewed some observational studies with strong biological hypotheses and adequate design allowing inferring causal links from intraspecific diversity to ecological dynamics (e.g. Post *et al.*, 2008). Studies varying intraspecific diversity within a set of multiple species (e.g. Booth & Grime, 2003) were not included in this meta-analysis. The literature search was carried out using the ISI *Web of Knowledge* and *Scopus* platforms (last accessed 25th July 2018). We also scrutinized the reference list of each article to obtain additional articles. The following key words were used in various combinations: *community genetics* AND *intraspecific variation, eco-evolutionary dynamics* AND *ecosystem function, community genetics* AND *ecosystem function*, and *intraspecific genetic variation* AND *ecosystem function*. We selected articles describing the effects of genotypic and/or phenotypic richness (intraspecific *richness*) and/or different genotypes/phenotypes (intraspecific *variation*) in a single target species on community and/or ecosystem dynamics. A total of 90 studies with available statistics were selected (see online Supporting information, Appendix S1 and Fig. S1). Among these, 23 studies (100% experimental studies) focused on intraspecific richness and 75 studies (90% experimental studies, 10% empirical studies) focused on intraspecific variation.

For each study, we recorded the Latin name of the target species and classified them as primary producers or consumers (including primary and secondary consumers as well as predators) and according to the major taxonomic categories represented in our data sets: arthropods (9 species), fishes (6 species), herbaceous plants (14 species), trees (31 species), and fungi (5 species). Overall, this led to 52 species for studies focusing on intraspecific variation, and 17 species for studies focusing on intraspecific richness. We recorded seven main response variables related to community structure and ecosystem functioning. A community is here defined as a group of at least two species, and we focused on three types of response variables describing the structure of communities: (1) species abundance: total number of individuals of all species; (2) biomass: total mass of individuals of all species; (3) community structure: number of species (e.g. Simpson or Shannon indices), species evenness and/or species richness.

Regarding response variables at the ecosystem level, we considered four main ecosystem functions: (1) decomposition rate: rate at which organic matter is recycled; (2) elemental cycling: quantity of organic or inorganic materials; (3) primary productivity: measured as several proxies of primary producers: biomass of primary producers excluding the productivity of the target species, chlorophyll *a* concentration, daily oxygen production, and photosynthetically active radiation; (4) ecosystem respiration: rate of oxygen consumption.

### (2) Meta-analysis

#### (a) The ecological consequences of intraspecific richness

To test for the consequences of intraspecific richness on ecological dynamics, we focused only on studies investigating the consequences of genotypic richness since this was the intraspecific diversity facet most commonly manipulated to test for the effects of intraspecific richness on ecological dynamics. Here, we used the log-transformed response ratio (ln*RR*) as an effect size. ln*RR* was computed as: ln 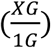, where *1G* is the average of the response variable measured for the treatment with a single genotype (i.e. monoculture), and *XG* is the average of the response variable measured for each treatment independently including more than one genotype. For each response variable, ln*RR* increases as the difference in the mean response variable measured in the treatment with a single genotype and treatments including more than one genotype increases. We also recorded the difference in the number of genotypes between the single genotype treatment (monoculture) and all other treatments separately (i.e. treatments including 2–12 genotypes) as the ‘difference in intraspecific richness’. In our data set, difference in intraspecific richness therefore varied between one and 11 genotypes. This approach allowed quantifying the ecological consequences of increasing the number of genotypes for each target species. Since each study generally assessed the effects of intraspecific richness on more than one response variable, our data set included a total of 135 assays.

We wanted to test the shape and the significance of the relationship between ln*RR* and the difference in intraspecific richness across all case studies. The general expectation is that ecological differences between treatments increase as differences in intraspecific richness increase, although this increase may be non-linear (Hughes *et al.*, 2008). We therefore used non-linear mixed-effect models to test the significance and shape of the relationship between absolute values of ln*RR* (|ln*RR*|) and differences in intraspecific richness. More precisely, we modelled this relationship using four different models to determine the most likely shape of the relationship between |ln*RR|* and difference in intraspecific richness: (1) a null model (one parameter) was computed for the null-effect hypothesis (i.e. no significant relationship between |ln*RR*| and difference in intraspecific richness); (2) a linear model (two parameters) suggesting a positive and linear relationship between |ln*RR*| and difference in intraspecific richness; (3) a Michaelis–Menten model (two parameters) in which |ln*RR*| increases with intraspecific richness, until a plateau is reached (saturating shape); (4) an asymptotic exponential model (two parameters) with a shape similar to the Michaelis–Menten model, except that the plateau is reached sooner.

All models cited above included article ID and the monoculture ID (i.e. the monoculture treatment to which each other treatment of richness was compared for a given response variable within each study) as random terms to account for non-independence of effect sizes (Noble *et al.*, 2017), and the inverse of the sample size as a weighting parameter giving greater weight to articles including more replicates. Models were compared using the Akaike information criterion (AIC) and we retained (as “best models”) all models that fell within a ΔAIC < 4 (Burnham & Anderson, 2002). We also calculated for each model the Akaike weight that provides a conditional probability for each model to be best supported by the data (Burnham & Anderson, 2002).

We then compared the magnitude (absolute effect size) of ecological effects of intraspecific and interspecific richness. We extracted from each study and for each response variable the ln*RR* corresponding to the most extreme levels of genotypic richness (*x*_max_) manipulated in each study (*N* = 63 ln*RR*). These values were subsequently compared to published ln*RR* values calculated following a similar method for experiments (*N* = 35) manipulating interspecific richness (Duffy *et al.*, 2017). Because absolute effect sizes follow a folded-normal distribution, we used an ‘analyse and transform’ approach (*sensu* Morrissey, 2016*a*,*b*) to estimate the absolute means of effect sizes. This approach consists first of estimating the mean and variance of ln*RR* (using non-absolute values), and then deriving the mean absolute value from these estimates. To do so, we estimated the mean of ln*RR* for interspecific and intraspecific richness, respectively, using two independent intercepts models with ln*RR* as the response variable, article ID as the random effect and the inverse of the sample size as the weighting parameter. These intercepts models were implemented using the MCMCglmm package in R (Hadfield, 2010). Markov chain Monte Carlo (MCMC) chains were run on 15×10^5^ iterations, with a burn-in interval of 3×10^4^, a thinning interval of 1×10^3^, and an inverse-Wishart prior (*V* = 1 and η = 0.002). Finally, the estimated means’ ln*RR* values were converted into absolute-magnitude |ln*RR*| values (following Morrissey, 2016*b*), and we compared the magnitudes of the ecological effects of interspecific and intraspecific richness based on visual inspection of 95% percentile intervals (PIs).

Finally, to compare the ecological consequences of intraspecific richness between levels of biological organization, we performed the same ‘analyse and transform’ approach described above. We used a linear mixed-effect model (implemented in the MCMCglmm package in R, and parameterized similarly than above) with the ln*RR* as the dependent variable, the inverse of the sample size as the weighting parameter, article ID and monoculture ID as random factors, and with the level of biological organization (community *versus* ecosystem response variables) treated as a fixed effect. The type of organism was not included in this analysis given that studies on intraspecific richness focused almost exclusively on primary producers (with two exceptions on fungi).

#### (b) The ecological consequences of intraspecific variation

Given that most studies (86%) did not include a control (i.e. a treatment without the target species), we compared the strength of effects among all unique genotypes and/or phenotypes that were considered in each study. Studies generally compared the consequences of 2–10 different unique genotypes and/or phenotypes (i.e. 2–10 treatments, with each treatment corresponding to a unique genotype/phenotype) on community and/or ecosystem dynamics; we gathered from each study the statistic (*t, F, Chi*-squared, Pearson’s *r*, Spearman’s *r, R*^*2*^ or Hedges’ *g*) associated with the between-treatments comparison (i.e. the variation of the phenotypes/genotypes). The higher the absolute value of the statistic, the higher the community and ecosystem consequences due to the variation of genotypes and/or phenotypes. The value of each statistic was converted into a correlation coefficient (*r*) ranging from 0 to 1 (see Table S1 for the formulae used). We did not use the direction of the statistic (i.e. positive or negative) because this depended upon the ecological response variable that was considered, which complicates comparisons on the direction of effects. The *Z*-Fisher transformation then was used to obtain a standardized effect size using the formula: 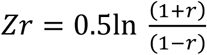. For each *Zr* value, we calculated the corresponding standard error as 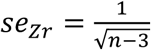 (Nakagawa & Cuthill, 2007). Since each study generally focused on more than one response variable, we obtained a total of 502 observed *Zr* values, each corresponding to the effect size of intraspecific variation observed within one species on a single response variable. The mean global *Zr* or mean effect size observed (MES_obs_ and its 95% confidence interval (CI) were calculated using an intercept-only model. This intercept-only model was run as a mixed model with no fixed effect, article ID as the random effect and *se*_*Zr*_ included as a weighting parameter to give more weight to studies with a larger sample size (Koricheva, Gurevitch & Mengersen, 2013).

Because *Zr* ranged between 0 and + ∞, the CIs of the MES_obs_ do not theoretically overlap 0, which makes it difficult to assess the significance of the strength of the MES_obs_. We therefore used a null-model approach to test if MESobs was significantly different from that expected under the null hypothesis, i.e. the true effect of intraspecific variation in all studies was zero. We resampled each statistic (e.g. *t, F*) from each empirical study in their respective null distribution with the adequate degree of freedom. This resampled set of statistics (*N* = 502) was transformed into *Zr* as described above to create a set of resampled *Zr* values. We used this set of resampled *Zr* values to fit an intercept-only model with no fixed effect, the corresponding article ID as a random term and *se*_zr_ included as a weighting parameter (as for MES_obs_). The mean global resampled *Zr* (MES_res_) was extracted from the model, and we repeated this resampling procedure 1000 times to obtain 1000 values of MES_res_. This resampled distribution of 1000 MES_res_ approximates the range of possible MES values expected if the null hypothesis was true. Finally, we calculated the probability of MES_obs_ to be larger than expected under this null hypothesis using a one-tailed test (Manly, 1997).

We then compared the median of effect sizes (MES_common_) of studies that were in common between our extended data set and that used by Des Roches *et al*. (2018) (i.e. 15 studies that were used both in our meta-analysis and that of Des Roches *et al*.) to a selection of 15 studies randomly sampled from our extended data set (i.e. 75 studies in our extended data set). We calculated the median effect size for the subset of random studies (MES_ran_) and repeated this resampling procedure 1000 times to obtain 1000 values of MESran. We then compared MES_common_ to each MES_ran_ value to calculate the probability that MES_common_ was higher than a random subset of 15 studies taken from the whole data set (Manly, 1997).

We then investigated the variability in effect sizes (*Zr*) and the potential moderators. We analysed the heterogeneity in effect sizes across articles using the *I*^2^ statistic, which was calculated using an intercept model with the article ID as the random effect and *se*_*Zr*_ as the weighting parameter (Higgins & Thompson, 2002; Senior *et al.*, 2016). Finally, we tested whether effect sizes (*Zr*) differed among organism types with intraspecific variation manipulation, and among the ecological response variables considered. We hence computed meta-regressions based on linear mixed-effect models with *Zr* values (for all 75 studies and 502 measures) as the dependent variable, and organism type or ecological response variable as fixed effects. The article ID was included as a random effect, and *se*_*Zr*_ was included as a weighting parameter. Four models were run to assess the differences of effect sizes (*i*) between organism types classified as consumers or primary producers, and (*ii*) between detailed taxonomic categories (arthropods, fishes, herbaceous plants and trees). We then tested whether the effect sizes of intraspecific variation differed among ecological response variables (*iii*) classified as community or ecosystem variables, and (*iv*) classified according to more detailed categories (abundance, biomass, community structure, decomposition, nutrient cycling, primary productivity and respiration of the ecosystem).

### (3) Publication bias

For both intraspecific variation and intraspecific richness, we assessed potential publication bias by combining Egger’s regressions and funnel plots (Egger *et al.*, 1997). Egger’s regressions and funnel plots were computed using the residuals of meta-regressions related effect sizes to the main modifiers (i.e. the explanatory variables) and a measure of study size (the inverse of *se*_*Zr*_ and sample size for intraspecific variation and intraspecific richness, respectively; Horvathova, Nakagawa & Uller, 2012; Nakagawa & Santos, 2012). Typically, for intraspecific variation we ran an Egger’s regression model including the residuals of the meta-regression linking intraspecific variation to the modifiers as a response variable and the inverse of *se*_*Zr*_ as the explanatory variable. A similar approach was used for intraspecific richness. The intercept α and the slope β of the Egger’s regressions are expected not to differ significantly from zero if the data sets are not biased towards significant results. Finally, funnel plots were produced as a scatterplot linking the residuals described above to the respective measure of the study size. An unbiased data set is expected to generate a funnel plot in which articles with larger sample sizes will be close to the mean effect size, whereas articles with small sample sizes will show more variance around the mean effect size (Horvathova *et al.*, 2012; Nakagawa & Santos, 2012).

Overall, and after accounting for important modifiers we found that there was no strong visual sign of publication bias, neither for intraspecific variation nor for intraspecific richness (Fig. S2). This visual inspection of funnel plots was confirmed by the Egger’s regressions since parameter values were not significant for intraspecific variation (α = 0.015, *P* = 0.404; β = –0.001, *P* = 0.501) or for intraspecific richness (α= –0.001, *P* = 0.914; β < 0.001, *P* = 0.961).

All statistical analyses were performed using the R environment (R core team, 2013, Appendix S2). The nlme package (Pinheiro *et al.*, 2014) was used to compute linear and non-linear mixed-effect models, unless specified otherwise.

## III. RESULTS

All articles (*N* = 90) selected for investigating the effects of intraspecific richness and intraspecific variation were published between 2000 and 2018, and 74% used primary producers as target species (Fig. 1). The first studies focusing on consumers were published in 2008, using fish (60%), arthropods (32%), and fungi (8%) as model species.

**Fig 1.**
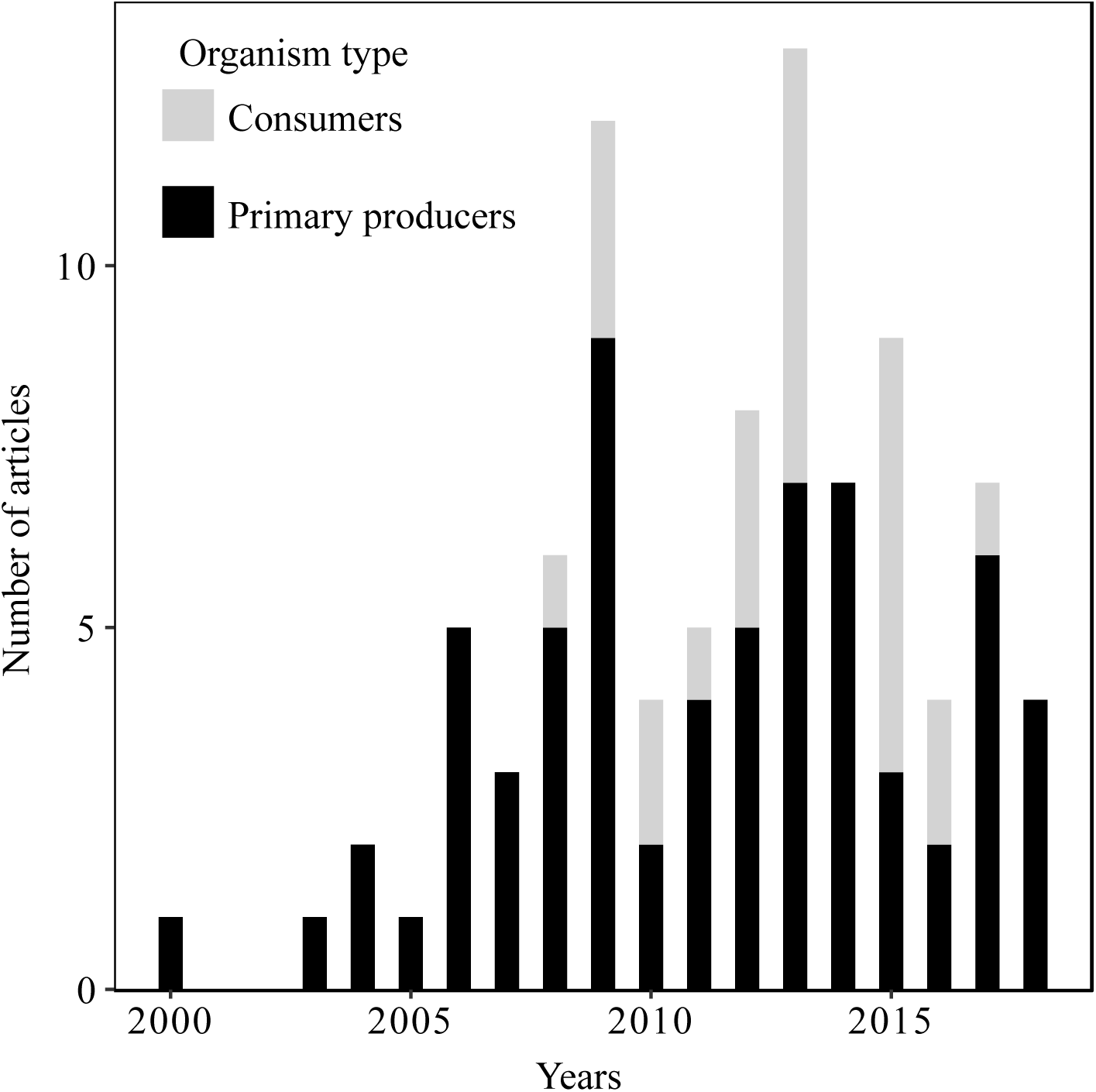
Publication year of the 90 selected articles used in the meta-analysis.

### (1) The intraspecific richness–ecological dynamics relationship

As expected, we found a significant, positive and non-linear relationship between intraspecific richness and ecological dynamics that approximated a saturating curve (Fig. 2A). The AIC selection procedure revealed that one out of the four tested models was highly likely to be supported by the data (i.e. 99% chance of being the best fitting model according to the Akaike weight and ΔAIC > 15.721 for the other models, Table 1). The model that best supported the data was the exponential asymptotic model, suggesting that the relationship between intraspecific richness and changes in community structure and ecosystem functioning (i.e. effect size: ln*RR*) likely followed a saturating shape (Fig. 2A).

**Table 1.** Summary table of model selection by Akaike information criterion (AIC) comparison to explain the shape of the relationship between the ecological consequences and the intraspecific richness. Models were run as non-linear mixed-effect models with the article ID as a random factor; equations and parameters estimates are also shown. IR, intraspecific richness; ln*RR*, effect size of intraspecific richness on ecological dynamics.

| Model | AIC | $\Delta AIC$ | AIC weight | Equation | Parameter estimates |
| --- | --- | --- | --- | --- | --- |
| Asymptotic exponential model | -176.695 | 0 | 0.999 | $\ln RR = a * (1 - e^{-b*IR})$ | $a = 0.221$<br>$b = 0.617$ |
| Michaelis–Menten model | -11.873 | 164.821 | < 0.001 | $\ln RR = V * \frac{IR}{k + IR}$ | $V = 0.054$<br>$k = -2.712$ |
| Linear model | -160.974 | 15.721 | < 0.001 | $\ln RR = b0 + b1 * IR$ | $b1 = 0.012$<br>$b0 = 0.122$ |
| Null model | -144.191 | 32.503 | < 0.001 | $\ln RR = b0$ | $b0 = 0.183$ |

**Fig 2.**
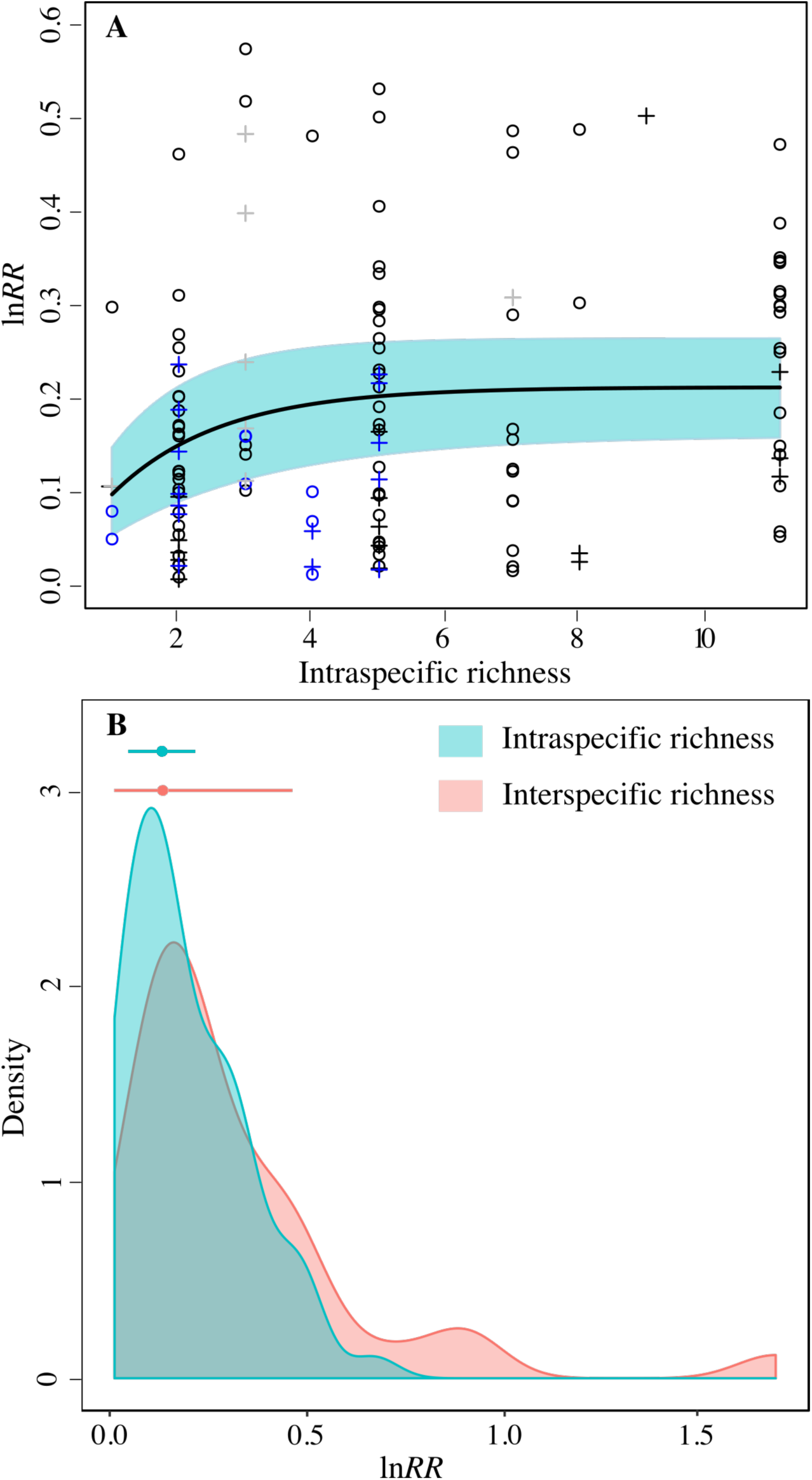
(A) Relationship between intraspecific richness and effect size (ln*RR*) on community (points) and ecosystem (crosses) dynamics. The line represents the shape of the relationship as predicted using an exponential asymptotic non-linear mixed effect model. The blue shadow represents 95% CI. Symbol colours denote the target species: herbaceous plant (black), tree (blue) or fungus (grey). (B) Density of absolute effect size (ln*RR*) for intraspecific and interspecific richness on ecological dynamics. Posterior means and 95% percentile intervals (points and horizontal lines, respectively) were estimated using a model including article ID as the random effect and the inverse of sample size as a weighting parameter.

We further found that the ecological effects of intraspecific richness were similar to the ecological effects induced by interspecific richness (Fig. 2B). Indeed, the two distributions largely overlapped and the estimated means were similar (intraspecific richness |ln*RR*| = 0.132, PI = 0.048–0.216; interspecific richness |ln*RR*| = 0.134 PI = 0.012–0.462). The ecological effects of intraspecific richness tended to be higher, although the difference was not significant, for community metrics (|ln*RR*| = 0.156, PI = 0.070–0.242) than for ecosystem metrics (|ln*RR*| = 0.045, PI = 0.004–0.137) (see Fig. S3 for details of ecological metrics).

### (2) The ecological consequences of intraspecific variation

We extended the meta-analysis performed by Des Roches *et al*. (2018) to 52 species (15 species were used in Des Roches *et al.*, 2018). We found that that the observed effect size of intraspecific variation on community structure and ecosystem dynamics was significant, and was twice as large as the resampled effect size expected under the null expectation (MES_obs_ = 0.521, 95% confidence interval (CI) = 0.444–0.598; MES_null_ = 0.259, CI = 0.258–0.259; resampled test, *P* < 0.001; see Figs 3 and S4).

**Fig 3.**
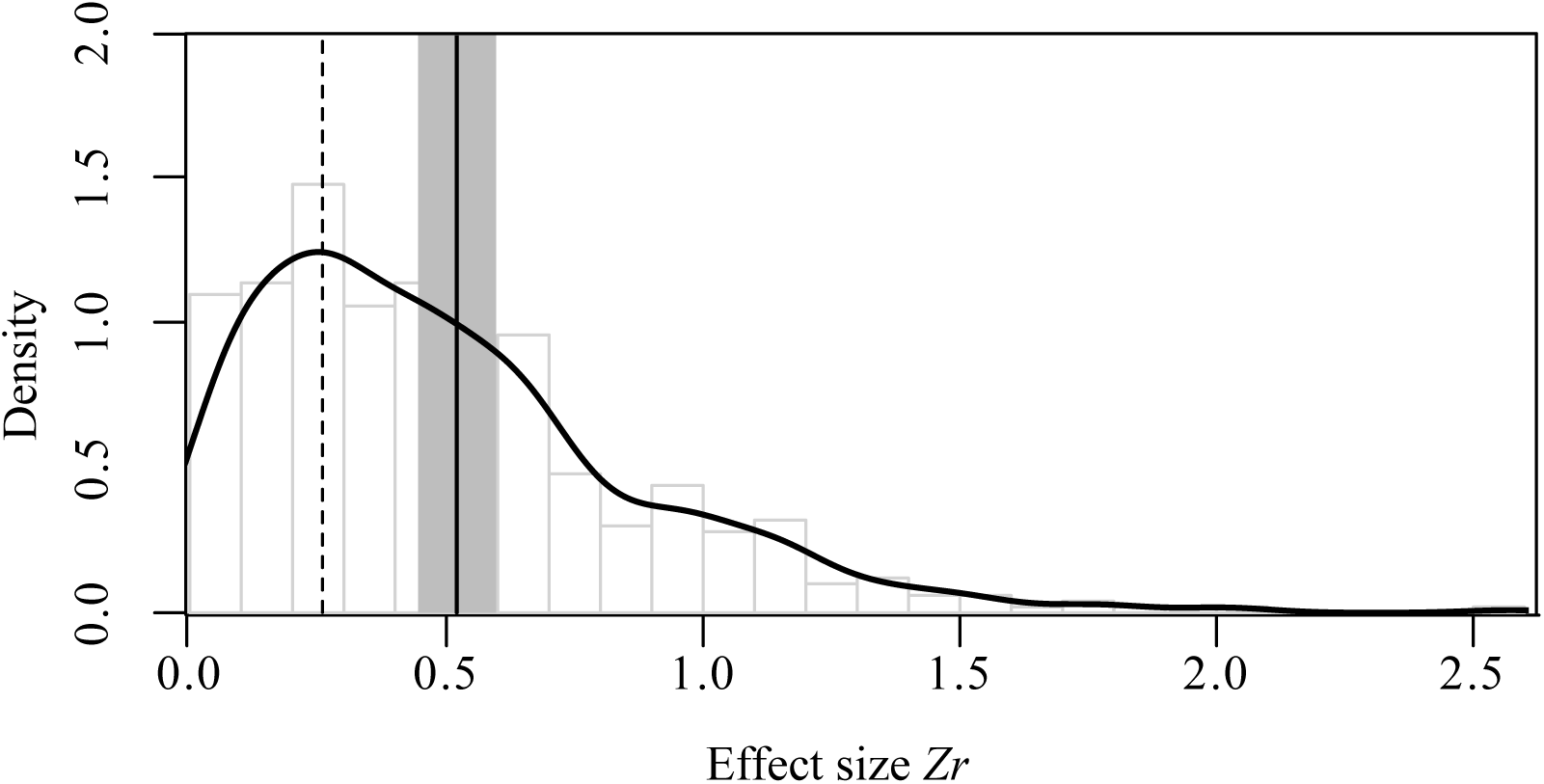
Density probability of effect size *Zr*. The vertical broken line represents the resampled *Zr* mean under the null hypothesis (confidence intervals not shown because they are too narrow); the black curve shows the distribution of observed *Zr* and its mean (vertical black straight line) and 95% CIs (grey shading).

We tested the extent to which the more restricted data set of Des Roches *et al.* (2018) was representative of our extended data set, or whether it was upwardly biased as expected by Des Roches *et al.* (2018). We found that that effect sizes for studies in common with the Des Roches *et al*. (2018) data set (MES_common_ = 0.299, 95% percentile interval (PI) = 0.033–1.092) were not significantly different from the distribution of effect sizes measured in our extended data set (MES_ran_ = 0.418, PI = 0.255–0.616; resampling test, *P* = 0.118; Fig. S5), and in fact showed a tendency to be downwardly biased.

Finally, a relatively low heterogeneity in effect size (*Zr*) was detected across articles (*I*^2^ = 0.151). The ecological effects induced by intraspecific variation were stronger when primary producers rather than consumers were manipulated (*F* = 3.968 d.f. = 1, 425, *P* = 0.047; Fig. 4A). Nonetheless, the strongest ecological effects of intraspecific variation tended to be observed in arthropods and herbaceous species, whereas the smallest effects were observed in fish and tree species (*F* = 2.475 d.f. = 3, 417, *P* = 0.061; Fig. 4A). Irrespective of organism type, the effects of intraspecific variation were significantly stronger when the response variables were measured at the ecosystem level rather than at the community level (*F* = 7.295, d.f = 1, 425, *P* = 0.007; Fig. 4B). The strongest effects were detected when response variables concerned nutrient cycling and the assembly of community, whereas the lowest effects were found for general measures of abundance and density of species (*F* = 2.725, d.f = 6, 417, *P* = 0.013; Fig. 4B).

**Fig 4.**
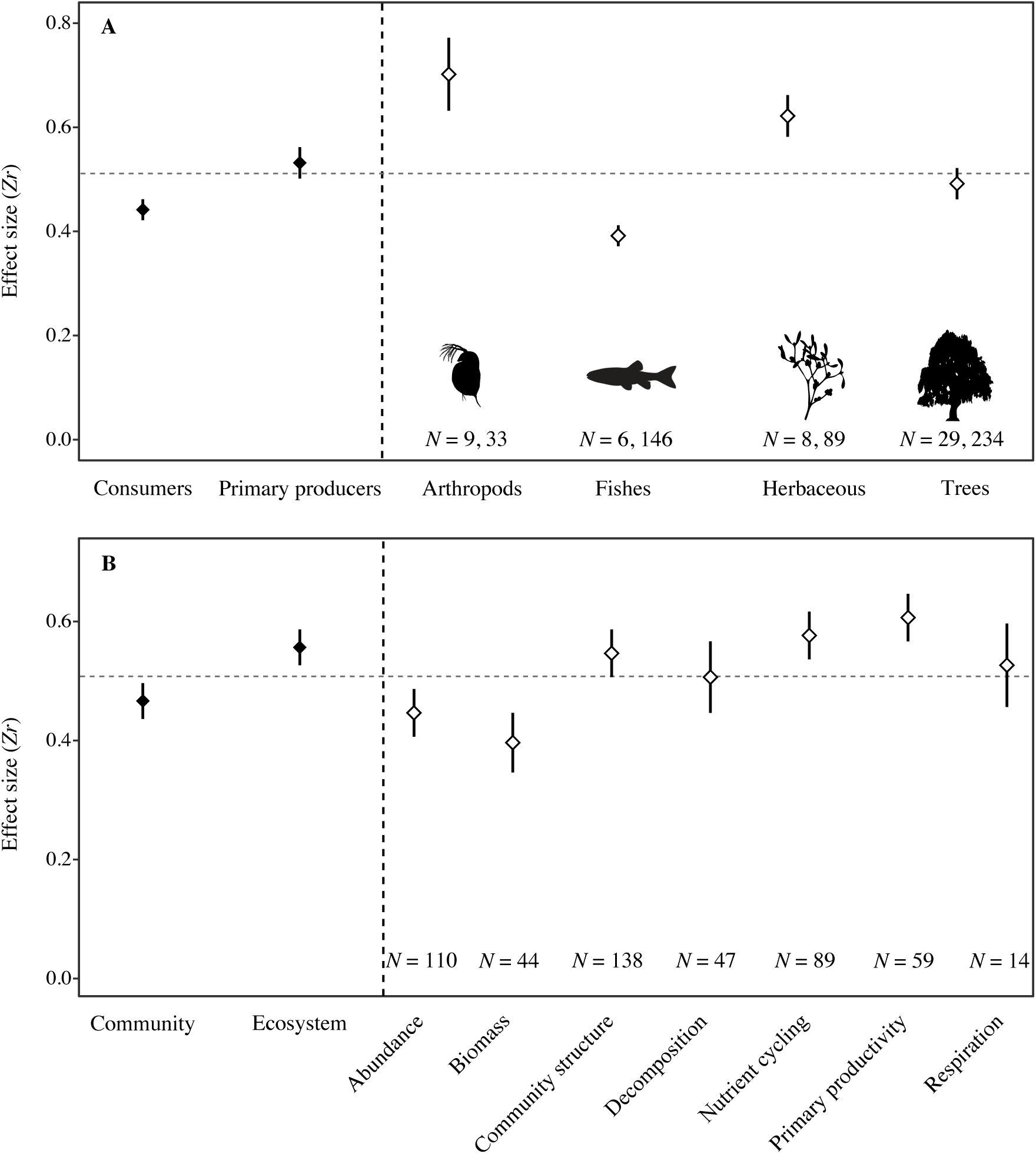
(A) Mean effect size *Zr* for different species groups. The sample sizes (*N*) represent the number of species and the number of effect sizes, respectively. The horizontal broken line represents the mean effect size; error bars represent ± 1 SE. (B) Mean *Zr* for the ecological response variables. The sample sizes (*N*) of the number of effect sizes are given. The horizontal broken line represents the mean effect size; error bars represent ± 1 SE.

## IV. DISCUSSION

Intraspecific diversity is increasingly recognized as an important facet of biodiversity that can affect all biological levels (Bailey *et al.*, 2009). Several studies have experimentally tested the ecological effects of intraspecific diversity, and we here provide the first global and quantitative estimates of the consequences of intraspecific richness and variation on community structure and ecosystem functioning. We demonstrated for the first time that the intraspecific BEF followed – as theoretically expected – a non-linear saturating curve with a plateau at 4–6 genotypes per population. Importantly, we demonstrated also for the first time that intraspecific richness affects community and ecosystem dynamics with a magnitude comparable to that of biodiversity measured at the species level. We further confirmed and extended the result that genotypic and/or phenotypic variation observed between populations can have non-negligible effects on community structure and ecosystem functions, and we demonstrated that previous estimates (Des Roches *et al.*, 2018) of these ecological effects of intraspecific variation actually tended to be underestimated. Finally, our exhaustive quantitative survey identified that the ecological consequences of intraspecific variation differ among biological level of organization, and among organism types. These findings provide novel and integrative insights, as well as multiple research perspectives, into the ecological role of intraspecific diversity.

### (1) Intraspecific diversity and the dynamics of communities and ecosystems

Although the form of the relationship between intraspecific richness and ecological consequences has already been discussed conceptually (Hughes *et al.*, 2008), our meta-analysis provides for the first time a qualitative and quantitative assessment of intraspecific BEF measured experimentally. Specifically, although considering mostly primary producers, our results demonstrated that an increase in intraspecific richness resulted in a non-linear (saturating) increase in the magnitude of its effects on ecological dynamics. This finding supports the idea that ecological divergence between an environment hosting populations composed of a single genotype and an environment hosting populations composed of multiple genotypes increases until a plateau is reached as the number of genotypes increases. This result echoes the BEF as defined at the interspecific level (Reiss *et al.*, 2009; Cadotte, Carscadden & Mirotchnick, 2011) and suggests that the saturating shape might arise from similar mechanisms occurring at the intraspecific and interspecific levels (Johnson *et al.*, 2006; Hughes *et al.*, 2008). More specifically, the initial linear increase is assumed to be due to complementarity and facilitation among genotypes, whereas the plateau likely occurs due to functional redundancy among genotypes (Johnson *et al.*, 2006). Redundant genotypes probably display functionally similar traits since two genotypes do not necessarily generate two functionally different traits (e.g. through synonymous mutations or trait convergences). Thus manipulating trait richness rather than genotypic richness, or more precisely manipulating functional effect traits [i.e. traits with ecological effects (Violle *et al.*, 2007)], in future experiments should allow us to explore the mechanisms underlying the intraspecific biodiversity–ecological dynamics relationship.

We found that effect sizes for intraspecific richness were very similar to values reported recently reported for experimental interspecific BEF, indicating that the ecological effects of varying phenotypic/genotypic richness within populations are close to those induced by varying species richness within communities. This finding raises several questions regarding the general relationships among intraspecific diversity, community structure, ecosystem functioning and common abiotic constraints. A large body of literature has demonstrated that intraspecific genetic diversity and species diversity (a measure of community structure) might co-vary because of common environmental drivers and/or reciprocal causal relationships between intraspecific genetic diversity and species diversity [i.e. the species–genetic diversity correlation (SGDC) framework (Vellend & Geber, 2005; Vellend, 2005)]. Because most studies considered in our meta-analysis are experimental, our findings confirm that intraspecific diversity can directly influence the structure of communities irrespective of the abiotic environments, hence adding weight to the SGDC framework. Additionally, we suggest expanding the SGDC framework since intraspecific diversity can also affect ecosystem functioning. This suggests that intraspecific diversity, community structure and ecosystem functioning may actually be tightly linked in a tripartite relationship. A major future challenge will be to tease apart the causal relationships linking these three components within a common abiotic environment. These relationships might be direct (e.g. intraspecific diversity directly affects community structure), indirect (e.g. intraspecific diversity indirectly affects ecosystem functions through its direct effect on community structure such as the trophic cascade), and/or due to the parallel effects of common abiotic drivers (e.g. temperature directly affects intraspecific diversity, community structure and ecosystem functions). As has been done recently for the BEF (Grace *et al.*, 2016; Duffy *et al.*, 2016) and the SGDC (Fourtune *et al.*, 2016; Lamy *et al.*, 2017) frameworks, we argue that a future important step will be to combine powerful statistical methods (e.g. path analysis; Shipley, 2000; Grace, 2006) with appropriate experimental designs to disentangle causal relationships between intraspecific diversity, community structure, ecosystem functions and their common environment.

We further demonstrated that intraspecific variation has significant ecological effects across a large set of species (52 species and 75 articles), hence confirming and refining a previous estimate based on a more restricted species set (Des Roches *et al.*, 2018). By more than tripling the number of species being investigated in this meta-analysis, we extend the conclusion to a greater taxonomic set that intraspecific variation is involved in shaping ecological dynamics, and that the ecological effects of intraspecific variation might be more common than expected. Moreover, we demonstrated that previous estimates (Des Roches *et al.*, 2018) were not upwardly biased (as expected from their focus on a non-random species pool), but were well within the range of estimates we report here and actually tended to be slight underestimates. Our finding hence strongly supports the idea that adaptive and non-adaptive processes can lead to unique populations differentially and significantly affecting ecological systems.

Although our conclusions held true for many species, the ecological effects of intraspecific variation were not homogeneous across species, and this was partly explained by their trophic level. Indeed, and according to expectations, the ecological effects of intraspecific variation were stronger when the target species was a primary producer than when it was a consumer. Several non-exclusive mechanisms might explain this result. For instance, many primary producers considered here provide a habitat for many invertebrate species (Southwood, Brown & Reader, 1979) (this is not the case for the consumer species), and this habitat can be modulated by changes in plant structure. The relative biomass of primary producers is higher than that of consumers, thus primary producers could generate stronger effects on communities and ecosystems than consumers simply because of this biomass effect. However, a more detailed analysis showed that the effects of intraspecific variation tended to be stronger for arthropod and herbaceous species than for fish (and to a lesser extent tree) species. This suggests that the trophic level of a species may not be the only predictor of the ecological effects of intraspecific diversity, and we argue that future work should aim to test specifically why intraspecific variation matters more for some species than others.

Finally, the effects of intraspecific variation were globally higher for ecosystems than for communities, hence generalizing across organism and ecosystem types a previous conclusion for freshwater consumer species (Palkovacs *et al.*, 2015). We can speculate that this difference arises because intraspecific variation acts on community dynamics through trophic mechanisms, whereas ecosystem functions can be modulated through both trophic and non-trophic interactions [e.g. excretion rate or leaf chemistry (Vanni, 2002; Schmitz *et al.*, 2014)]. For instance, a consumer species that shows intraspecific variation in resource selectivity and/or consumption rate could affect both the community structure and productivity of its resource (Harmon *et al.*, 2009). Non-trophic mechanisms such as variability in organismal stoichiometry could reinforce the effect of the consumer species on several ecosystem functions [e.g. primary production or soil mineralization (Schmitz, Hawlena & Trussell, 2010; Hawlena *et al.*, 2012)]. Alternatively, changes in ecosystem functions might be due to both direct effects of intraspecific diversity and indirect effects of intraspecific diversity mediated through changes in community structure, which may overall strengthen the effects of intraspecific diversity at the ecosystem level. However, a more detailed analysis revealed that much variation exists between sub-categories of response variables (Fig. 4B), and that the dichotomy between variables measured at the community and ecosystem levels is not straightforward. Although providing the first attempts to separate variance in the ecological effects of intraspecific variation into its component parts, our findings call for further studies on various taxa and in different ecosystems in order to understand fully the ecological effects of intraspecific diversity.

### (2) Research perspectives on the relationships between intraspecific diversity and the dynamics of communities and ecosystems

We highlight several research avenues that may greatly enhance our understanding of the relationships between intraspecific diversity and the dynamics of communities and ecosystems.

Our review demonstrates that the ecological effect size of intraspecific diversity varies among species and that this interspecific variance in effect size can be partly explained by the type of organism (i.e. primary producer or consumer). However, species composing a community also vary according to abundance, role in the ecosystem (e.g. keystone species; Paine, 1969), body size (or height for plants), life-history strategy (e.g. *r–K* strategy), recent history (e.g. whether the species is non-native), functional characteristics (e.g. stoichiometry, metabolism), etc. The next challenge will be to partition this interspecific variance in effect size better by determining the species characteristics that best predict the strength of effect sizes; this is a pre-requisite to design coherent conservation plans at the intraspecific level (Mimura *et al.*, 2016).

Intraspecific diversity is often manipulated with respect to a single target species. However, in nature, species are interacting and we argue that future studies should manipulate intraspecific diversity within multiple interacting species to reflect natural conditions, and to allow partitioning the relative importance of intraspecific diversity between interacting (and potentially co-evolving) species.

Ecosystems are interconnected through cross-ecosystem fluxes of matter (Loreau, Mouquet & Holt, 2003). For instance, freshwater ecosystems such as rivers receive a large amount of material from surrounding terrestrial ecosystems [e.g. dead leaves falling from trees (Bartels *et al.*, 2012)]. A major future challenge would be to assess the relative effects of intraspecific diversity on allochtonous ecosystems *versus* autochtonous ecosystems; for example testing whether the consequences of intraspecific diversity manipulation in a fish species are greater in associated aquatic ecosystems than on nearby terrestrial ecosystems [see Jackrel & Wootton (2014), Crutsinger *et al.* (2014)*b* and Rudman *et al.* (2015) for attempts to quantify across-ecosystem effects of intraspecific diversity]. This is an important next step to quantify in a more integrative way the importance of intraspecific diversity in natural systems.

Previous studies have mainly assessed the consequences of intraspecific diversity by considering the genetic or the phenotypic characteristics of organisms as a whole. However, some traits have been shown to be more important for ecological dynamics than others; this is the case for functional effect traits that are defined as traits with an ecological impact [e.g. excretion rate or leaf nutrient content (Violle *et al.*, 2007; Díaz *et al.*, 2013)]. We suggest that direct manipulation of the functional trait diversity of individuals within a species, rather than their genotypic or phenotypic diversity as a whole would be a powerful means to understanding the mechanisms by which intraspecific diversity acts on ecological dynamics.

Although we emphasize that intraspecific diversity is an essential component of the community and ecosystem facets of ecological dynamics, comparisons with the effects of other major ecological constraints (e.g. temperature, interspecific biodiversity, human disturbance) have rarely been conducted (but see Burkle *et al.*, 2013; El-Sabaawi *et al.*, 2015). To evaluate and quantify the importance of intraspecific diversity in natural systems better, it is important to assess the relative contributions of intraspecific diversity and other determinants of ecosystem functioning. This will be an essential step in confirming that intraspecific variation is a key determinant, and not just a random signal in complex systems.

## V. CONCLUSIONS

(1) This study provides empirical evidence that the ecological effects of intraspecific richness increase asymptotically, paralleling well-known patterns observed at the interspecific level (Loreau, 2000; Hooper *et al.*, 2005) and confirming previous hypotheses (Hughes *et al.*, 2008).

(2) We found that experimental manipulations of intraspecific richness caused community and ecosystem differentiations as large as those generated by interspecific richness. This suggests that variation in richness within populations can have similar ecological consequences to variation in richness among species.

(3) We also demonstrated that variation in phenotypes or genotypes within species is an important driver of community and ecosystem dynamics. These major ecological effects of intraspecific diversity held true for a range of organisms including plants and animals, although much remains to be tested.

(4) Overall these findings demonstrate that intraspecific diversity - beyond its importance for species to adapt to environmental changes - is an important facet of biodiversity for understanding and predicting the ecological dynamics of communities and ecosystems, reinforcing the need for a better appraisal of the causes and consequences of intraspecific diversity in natural populations and for improved conservation plans.

## VI. ACKNOWLEDGEMENTS

We warmly thank José M. Montoya for valuable comments on a previous version of this review. We also thank three anonymous reviewers for their constructive and useful comments. A.R. was supported financially by a Doctoral scholarship from the Université Fédérale de Toulouse. This work was undertaken at SETE, which is part of the “Laboratoire d’Excellence” (LABEX) entitled TULIP (ANR-10-LABX-41).

## VIII. SUPPORTING INFORMATION

Additional supporting information may be found in the online version of this article.

**Appendix S1.** Papers used in the meta-analysis.

**Fig. S1.** Flow diagram of manuscript screening and eligibility for this meta-analysis.

**Table S1.** Formulae used to convert different statistical values into an *r* correlation coefficient.

**Fig. S2.** Funnel plots describing the residuals effect size distribution against an estimate of the study size for (A) intraspecific variation and (B) intraspecific richness.

**Appendix S2.** R script of the principal functions used in analyses.

**Fig, S3.** Distribution of effect sizes (ln*RR*) across ecological metrics.

**Fig. S4.** Distribution of resampled mean effect size (MES) expected under the null hypothesis, i.e. the effect of intraspecific identity in all studies is zero.

**Fig. S5.** Distribution of median effect sizes calculated for each of the 1000 sets of 15 studies randomly selected within our extended data set of 75 studies.

## REFERENCES

Abbott, J.M., Grosberg, R.K., Williams, S.L. & Stachowicz, J.J. (2017). Multiple dimensions of intraspecific diversity affect biomass of eelgrass and its associated community. Ecology 98, 3152–3164.

Bailey, J.K., Schweitzer, J.A., Ubeda, F., Koricheva, J., Leroy, C.J., Madritch, M.D., Rehill, B.J., Bangert, R.K., Fischer, D.G., Allan, G.J. & Whitham, T.G. (2009). From genes to ecosystems: a synthesis of the effects of plant genetic factors across levels of organization. Philosophical Transactions of the Royal Society B: Biological Sciences 364, 1607–1616.

Barbour, M.A., Fortuna, M.A., Bascompte, J., Nicholson, J.R., Julkunen-Tiitto, R., Jules, E.S. & Crutsinger, G.M. (2016). Genetic specificity of a plant–insect food web: Implications for linking genetic variation to network complexity. Proceedings of the National Academy of Sciences 113, 2128–2133.

Barbour, R.C., Forster, L.G., Baker, S.C., Steane, D.A. & Potts, B.M. (2009a). Biodiversity consequences of genetic variation in bark characteristics within a foundation tree species. Conservation Biology 23, 1146–1155.

Barbour, R.C., O’Reilly-Wapstra, J.M., De Little, D.W., Jordan, G.J., Steane, D.A., Humphreys, J.R., Bailey, J.K., Whitham, T.G. & Potts, B.M. (2009b). A geographic mosaic of genetic variation within a foundation tree species and its community-level consequences. Ecology 90, 1762–1772.

Barbour, R.C., Storer, M.J. & Potts, B.M. (2009c). Relative importance of tree genetics and microhabitat on macrofungal biodiversity on coarse woody debris. Oecologia 160, 335–342.

Barrios-Garcia, M.N., Rodriguez-Cabal, M.A., Rudgers, J.A. & Crutsinger, G.M. (2016). Soil fertilization does not alter plant architectural effects on arthropod communities. Journal of Plant Ecology, rtw087.

Bartels, P., Cucherousset, J., Steger, K., Eklöv, P., Tranvik, L.J. & Hillebrand, H (2012). Reciprocal subsidies between freshwater and terrestrial ecosystems structure consumer resource dynamics. Ecology 93, 1173–1182.

Bassar, R.D., Marshall, M.C., Lopez-Sepulcre, A., Zandona, E., Auer, S.K., Travis, J., Pringle, C.M., Flecker, A.S., Thomas, S.A., Fraser, D.F. & Reznick, D.N. (2010). Local adaptation in Trinidadian guppies alters ecosystem processes. Proceedings of the National Academy of Sciences 107, 3616–3621.

Bolnick, D.I., Amarasekare, P., Araújo, M.S., Bürger, R., Levine, J.M., Novak, M., Rudolf, V.H., Schreiber, S.J. & Urban, M.C. (2011). Why intraspecific trait variation matters in community ecology. Trends in ecology & evolution 26, 183–192.

Bolnick, D.I., Svanbäck, R., Fordyce, J.A., Yang, L.H., Davis, J.M., Hulsey, C.D. & Forister, M.L. (2003). The ecology of individuals: incidence and implications of individual specialization. The American Naturalist 161, 1–28.

Booth, R.E. & Grime, J.P. (2003). Effects of genetic impoverishment on plant community diversity. Journal of Ecology 91, 721–730.

Burkle, L.A., Souza, L., Genung, M.A. & Crutsinger, G.M. (2013). Plant genotype, nutrients, and G X E interactions structure floral visitor communities. Ecosphere 4.

Burnham, K.P. & Anderson, D.R. (2002). Model selection and multimodel inference: a practical information-theoretic approach. Springer, New York.

Bustos-Segura, C., Poelman, E.H., Reichelt, M., Gershenzon, J. & Gols, R. (2017). Intraspecific chemical diversity among neighbouring plants correlates positively with plant size and herbivore load but negatively with herbivore damage. Ecology Letters 20, 87–97.

Cadotte, M.W., Carscadden, K. & Mirotchnick, N. (2011). Beyond species: functional diversity and the maintenance of ecological processes and services. Journal of Applied Ecology 48, 1079–1087.

Campos-Navarrete, M.J., Munguía-Rosas, M.A., Abdala-Roberts, L., Quinto, J. & Parra-Tabla, V. (2015). Effects of tree genotypic diversity and species diversity on the arthropod community associated with big-leaf mahogany. Biotropica 47, 579–587.

Cardinale, B.J., Duffy, J.E., Gonzalez, A., Hooper, D.U., Perrings, C., Venail, P., Narwani, A., Mace, G.M., Tilman, D., Wardle, D.A., Kinzig, A.P., Daily, G.C., Loreau, M., Grace, J.B., Larigauderie, A., ET AL. (2012). Biodiversity loss and its impact on humanity. Nature 486, 59–67.

Chapin, S.F., Zavaleta, E.S., Eviner, V.T., Naylor, R.L., Vitousek, P.M., Reynolds, H.L., Hooper, D.U., Lavorel, S., Sala, O.E., Hobbie, S.E., Mack, M.C. & Díaz, S. (2000). Consequences of changing biodiversity. Nature 405, 234–242.

Chislock, M.F., Sarnelle, O., Olsen, B.K., Doster, E. & Wilson, A.E. (2013). Large effects of consumer offense on ecosystem structure and function. Ecology 94, 2375–2380.

Classen, A.T., Chapman, S.K., Whitham, T.G., Hart, S.C. & Koch, G.W. (2007). Genetic-based plant resistance and susceptibility traits to herbivory influence needle and root litter nutrient dynamics. Journal of Ecology 95, 1181–1194.

Cook-Patton, S.C., Mcart, S.H., Parachnowitsch, A.L., Thaler, J.S. & Agrawal, A.A. (2011). A direct comparison of the consequences of plant genotypic and species diversity on communities and ecosystem function. Ecology 92, 915–923.

Cornelissen, J.H.C., PEREZ-Harguindeguy, N., Gwynn-Jones, D., Diaz, S., Callaghan, T.V. & Aerts, R. (2000). Autumn leaf colours as indicators of decomposition rate in sycamore (*Acer pseudoplatanus L*.). Plant and Soil 225, 33–38.

Crawford, K.M. & Rudgers, J.A. (2013). Genetic diversity within a dominant plant outweighs plant species diversity in structuring an arthropod community. Ecology 94, 1025–1035.

Crutsinger, G.M., Cadotte, M.W. & Sanders, N.J. (2009a). Plant genetics shapes inquiline community structure across spatial scales. Ecology Letters 12, 285–292.

Crutsinger, G.M., Carter, B.E. & Rudgers, J.A. (2013). Soil nutrients trump intraspecific effects on understory plant communities. Oecologia 173, 1531–1538.

Crutsinger, G.M., Collins, M.D., Fordyce, J.A., Gompert, Z., Nice, C.C. & Sanders, N.J. (2006). Plant genotypic diversity predicts community structure and governs an ecosystem process. Science 313, 966–968.

Crutsinger, G.M., Collins, M.D., Fordyce, J.A. & Sanders, N.J. (2008a). Temporal dynamics in non-additive responses of arthropods to host-plant genotypic diversity. Oikos 117, 255–264.

Crutsinger, G.M., Reynolds, W.N., Classen, A.T. & Sanders, N.J. (2008b). Disparate effects of plant genotypic diversity on foliage and litter arthropod communities. Oecologia 158, 65–75.

Crutsinger, G.M., Rodriguez-Cabal, M.A., Roddy, A.B., Peay, K.G., Bastow, J.L., Kidder, A.G., Dawson, T.E., Fine, P.V.A. & Rudgers, J.A. (2014a).Genetic variation within a dominant shrub structures green and brown community assemblages. Ecology 95, 387–398.

Crutsinger, G.M., Rudman, S.M., Rodriguez-Cabal, M.A., Mckown, A.D., Sato, T., Macdonald, A.M., Heavyside, J., Geraldes, A., Hart, E.M., Leroy, C.J. & El-Sabaawi, R.W. (2014b). Testing a ‘genes-to-ecosystems’ approach to understanding aquatic-terrestrial linkages. Molecular Ecology 23, 5888–5903.

Crutsinger, G.M., Sanders, N.J. & Classen, A.T. (2009b). Comparing intra- and inter-specific effects on litter decomposition in an old-field ecosystem. Basic and Applied Ecology 10, 535–543.

De Bello, F., Lavorel, S., Albert, C.H., Thuiller, W., Grigulis, K., Dolezal, J., Janecek, S. & Leps, J. (2011). Quantifying the relevance of intraspecific trait variability for functional diversity. Methods in Ecology and Evolution 2, 163–174.

De Graaff, M.-A., Six, J., Jastrow, J.D., Schadt, C.W. & Wullschleger, S.D. (2013). Variation in root architecture among switchgrass cultivars impacts root decomposition rates. Soil Biology & Biochemistry 58, 198–206.

Des Roches, S., Post, D.M., Turley, N.E., Bailey, J.K., Hendry, A.P., Kinnison, M.T., Schweitzer, J.A. & Palkovacs, E.P. (2018). The ecological importance of intraspecific variation. Nature Ecology & Evolution 2, 57–64.

Des Roches, S., Shurin, J.B., Schluter, D. & Harmon, L.J. (2013). Ecological and evolutionary effects of stickleback on community structure. PLoS ONE 8, e59644.

Díaz, S., Purvis, A., Cornelissen, J.H.C., Mace, G.M., Donoghue, M.J., Ewers, R.M., Jordano, P. & Pearse, W.D. (2013). Functional traits, the phylogeny of function, and ecosystem service vulnerability. Ecology and Evolution 3, 2958–2975.

Downing, A.L. & Leibold, M.A. (2002). Ecosystem consequences of species richness and composition in pond food web. Nature 416, 837–841.

Duffy, J.E., Godwin, C.M. & Cardinale, B.J. (2017). Biodiversity effects in the wild are common and as strong as key drivers of productivity. Nature 549, 261–264.

Duffy, J.E., Lefcheck, J.S., Stuart-Smith, R.D., Navarrete, S.A. & Edgar, G.J. (2016). Biodiversity enhances reef fish biomass and resistance to climate change. Proceedings of the National Academy of Sciences 113, 6230–6235.

Egger, M., Smith, G.D., Schneider, M. & Minder, C. (1997). Bias in meta-analysis detected by a simple graphical test. Bmj 315, 629–634.

El-Sabaawi, R.W., Bassar, R.D., Rakowski, C., Marshall, M.C., Bryan, B.L., Thomas, S.N., Pringle, C., Reznick, D.N. & Flecker, A.S. (2015). Intraspecific phenotypic differences in fish affect ecosystem processes as much as bottom-up factors. Oikos 124, 1181–1191.

Farkas, T.E., Mononen, T., Comeault, A.A., Hanski, I. & Nosil, P. (2013). Evolution of camouflage drives rapid ecological change in an insect community. Current Biology 23, 1835–1843.

Fourtune, L., Paz-Vinas, I., Loot, G., Prunier, J.G. & Blanchet, S. (2016). Lessons from the fish: a multi-species analysis reveals common processes underlying similar species-genetic diversity correlations. Freshwater Biology 61, 1830–1845.

Fridley, J.D. & Grime, J.P. (2010). Community and ecosystem effects of intraspecific genetic diversity in grassland microcosms of varying species diversity. Ecology 91, 2272–2283.

Fryxell, D.C., Arnett, H.A., Apgar, T.M., Kinnison, M.T. & Palkovacs, E.P. (2015). Sex ratio variation shapes the ecological effects of a globally introduced freshwater fish. Proceedings of the Royal Society B: Biological Sciences 282, 20151970.

Genung, M.A., Lessard, J.-P., Brown, C.B., Bunn, W.A., Cregger, M.A., Reynolds, W.N., Felker-Quinn, E., Stevenson, M.L., Hartley, A.S., Crutsinger, G.M., Schweitzer, J.A. & Bailey, J.K. (2010). Non-additive effects of genotypic diversity increase floral abundance and abundance of floral visitors. Plos One 5, e8711.

Goitom, E., Kilsdonk, L.J., Brans, K., Jansen, M., Lemmens, P. & De Meester, L. (2018). Rapid evolution leads to differential population dynamics and top-down control in resurrected *Daphnia* populations. Evolutionary Applications 11, 96–111.

Grace, J.B. (2006). Structural equation modeling and natural systems. Cambridge University Press.

Grace, J.B., Anderson, T.M., Seabloom, E.W., Borer, E.T., Adler, P.B., Harpole, W.S., Hautier, Y., Hillebrand, H., Lind, E.M., Pärtel, M., Bakker, J.D., Buckley, Y.M., Crawley, M.J., Damschen, E.I., Davies, K.F., ET AL. (2016). Integrative modelling reveals mechanisms linking productivity and plant species richness. Nature 529, 390–393.

Grant, P.R. & Grant, B.R. (2006). Evolution of character displacement in darwin’s finches. Science 313, 224–226.

Hadfield, J.D. (2010). MCMC methods for multi-response generalized linear mixed models: the MCMCglmm R package. Journal of Statistical Software 33, 1–22.

Harmon, L.J., Matthews, B., Des Roches, S., Chase, J.M., Shurin, J.B. & Schluter, D. (2009). Evolutionary diversification in stickleback affects ecosystem functioning. Nature 458, 1167–1170.

Harrison, J.G., Philbin, C.S., Gompert, Z., Forister, G.W., Hernandez-Espinoza, L., Sullivan, B.W., Wallace, I.S., Beltran, L., Dodson, C.D., Francis, J.S., Schlageter, A., Shelef, O., Yoon, S.A. & Forister, M.L. (2018). Deconstruction of a plant-arthropod community reveals influential plant traits with nonlinear effects on arthropod assemblages. Functional Ecology 32, 1317–1328.

Hawlena, D., Strickland, M.S., Bradford, M.A. & Schmitz, O.J. (2012). Fear of predation slows plant-litter decomposition. Science 336, 1434–1438.

Higgins, J.P.T. & Thompson, S.G. (2002). Quantifying heterogeneity in a meta-analysis. Statistics in Medicine 21, 1539–1558.

Hillebrand, H. & Matthiessen, B. (2009). Biodiversity in a complex world: consolidation and progress in functional biodiversity research. Ecology Letters 12, 1405–1419.

Hines, J., Reyes, M., Mozder, T.J. & Gessner, M.O. (2014). Genotypic trait variation modifies effects of climate warming and nitrogen deposition on litter mass loss and microbial respiration. Global Change Biology 20, 3780–3789.

Hooper, D.U., Chapin, F.S., Ewel, J.J., Hector, A., Inchausti, P., Lavorel, S., Lawton, J.H., Lodge, D.M., Loreau, M., Naeem, S., Schmid, B., Setälä, H., Symstad, A.J., Vandermeer, J. & Wardle, D.A. (2005). Effects of biodiversity on ecosystem functioning: a consensus of current knowledge. Ecological Monographs 75, 3–35.

Horvathova, T., Nakagawa, S. & Uller, T. (2012). Strategic female reproductive investment in response to male attractiveness in birds. Proceedings of the Royal Society B: Biological Sciences 279, 163–170.

Howeth, J.G., Weis, J.J., Brodersen, J., Hatton, E.C. & Post, D.M. (2013). Intraspecific phenotypic variation in a fish predator affects multitrophic lake metacommunity structure. Ecology and Evolution 3, 5031–5044.

Huang, J., Liu, M., Chen, X., Chen, J., Li, H. & Hu, F. (2015). Effects of intraspecific variation in rice resistance to aboveground herbivore, brown planthopper, and rice root nematodes on plant yield, labile pools of plant, and rhizosphere soil. Biology and Fertility of Soils 51, 417–425.

Hughes, A.R. (2014). Genotypic diversity and trait variance interact to affect marsh plant performance. Journal of Ecology 102, 651–658.

Hughes, A.R., Inouye, B.D., Johnson, M.T.J., Underwood, N. & Vellend, M. (2008). Ecological consequences of genetic diversity. Ecology Letters 11, 609–623.

Ingram, T., Svanbäck, R., Kraft, N.J.B., Kratina, P., Southcott, L. & Schluter, D. (2012). Intraguild predation drives evolutionary niche shift in threespine stickleback. Evolution 66, 1819–1832.

Ito, M. & Ozaki, K. (2005). Response of a gall wasp community to genetic variation in the host plant *Quercus crispula*: a test using half-sib families. Acta Oecologica 27, 17–24.

Jackrel, S.L. & Wootton, J.T. (2014). Local adaptation of stream communities to intraspecific variation in a terrestrial ecosystem subsidy. Ecology 95, 37–43.

Johnson, M.T. & Agrawal, A.A. (2005). Plant genotype and environment interact to shape a diverse arthropod community on evening primrose (*Oenothera biennis*). Ecology 86, 874–885.

Johnson, M.T.J., Lajeunesse, M.J. & Agrawal, A.A. (2006). Additive and interactive effects of plant genotypic diversity on arthropod communities and plant fitness. Ecology Letters 9, 24–34.

Johnson, M.T.J., Vellend, M. & Stinchcombe, J.R. (2009). Evolution in plant populations as a driver of ecological changes in arthropod communities. Philosophical Transactions of the Royal Society B: Biological Sciences 364, 1593–1605.

Kagiya, S., Yasugi, M., Kudoh, H., Nagano, A.J. & Utsumi, S. (2018). Does genomic variation in a foundation species predict arthropod community structure in a riparian forest? Molecular Ecology 27, 1284–1295.

Kanaga, M.K., Latta, L.C., Mock, K.E., Ryel, R.J., Lindroth, R.L. & Pfrender, M.E. (2009). Plant genotypic diversity and environmental stress interact to negatively affect arthropod community diversity. Arthropod-Plant Interactions 3, 249–258.

Katano, O. (2011). Effects of individual differences in foraging of pale chub on algal biomass through trophic cascades. Environmental Biology of Fishes 92, 101–112.

Koricheva, J., Gurevitch, J. & Mengersen, K. (2013). Handbook of meta-Analysis in ecology and evolution. Princeton University press, Princeton, New Jersey.

Kotowska, A.M., Cahill JR, J.F. & Keddie, B.A. (2010). Plant genetic diversity yields increased plant productivity and herbivore performance. Journal of Ecology 98, 237–245.

Lagerstrom, A., Nilsson, M.-C. & Wardle, D.A. (2013). Decoupled responses of tree and shrub leaf and litter trait values to ecosystem retrogression across an island area gradient. Plant and Soil 367, 183–197.

Lamy, T., Laroche, F., David, P., Massol, F. & Jarne, P. (2017). The contribution of species-genetic diversity correlations to the understanding of community assembly rules. Oikos 126, 759–771.

Lecerf, A. & Chauvet, E. (2008). Intraspecific variability in leaf traits strongly affects alder leaf decomposition in a stream. Basic and Applied Ecology 9, 598–605.

Leroy, C.J., Whitham, T.G., Wooley, S.C. & Marks, J.C. (2007). Within-species variation in foliar chemistry influences leaf-litter decomposition in a Utah river. Journal of the North American Benthological Society 26, 426–438.

Loreau, M. (2000). Biodiversity and ecosystem functioning: recent theoretical advances. Oikos 91, 3–17.

Loreau, M., Mouquet, N. & Holt, R.D. (2003). Meta-ecosystems: a theoretical framework for a spatial ecosystem ecology. Ecology Letters 6, 673–679.

Lowe, W.H., Kovach, R.P. & Allendorf, F.W. (2017). Population genetics and demography unite ecology and evolution. Trends in Ecology & Evolution 32, 141–152.

Madritch, M., Donaldson, J.R. & Lindroth, R.L. (2006). Genetic identity of *Populus tremuloides* litter influences decomposition and nutrient release in a mixed forest stand. Ecosystems 9, 528–537.

Madritch, M.D., Greene, S.L. & Lindroth, R.L. (2009). Genetic mosaics of ecosystem functioning across aspen-dominated landscapes. Oecologia 160, 119–127.

Madritch, M.D. & Hunter, M.D. (2003). Intraspecific litter diversity and nitrogen deposition affect nutrient dynamics and soil respiration. Oecologia 136, 124–128.

Madritch, M.D. & Hunter, M.D. (2004). Phenotypic diversity and litter chemistry affect nutrient dynamics during litter decomposition in a two species mix. Oikos 105, 125–131.

Madritch, M.D. & Hunter, M.D. (2005). Phenotypic variation in oak litter influences short- and long-term nutrient cycling through litter chemistry. Soil Biology & Biochemistry 37, 319–327.

Madritch, M.D. & Lindroth, R.L. (2011). Soil microbial communities adapt to genetic variation in leaf litter inputs. Oikos 120, 1696–1704.

Manly, B.F. (1997). Randomization, bootstrap and Monte Carlo methods in biology. Chapman and Hall, London.

Matthews, B., Aebischer, T., Sullam, K.E., Lundsgaard-Hansen, B. & Seehausen, O. (2016). Experimental evidence of an eco-evolutionary feedback during adaptive divergence. Current Biology 26, 483–489.

Matthews, B., De Meester, L., Jones, C.G., Ibelings, B.W., Bouma, T.J., Nuutinen, V., De Koppel, J. VAN & Odling-Smee, J. (2014). Under niche construction: an operational bridge between ecology, evolution, and ecosystem science. Ecological Monographs 84, 245–263.

Matthews, B., Narwani, A., Hausch, S., Nonaka, E., Peter, H., Yamamichi, M., Sullam, K.E., Bird, K.C., Thomas, M.K., Hanley, T.C. & Turner, C.B. (2011). Toward an integration of evolutionary biology and ecosystem science. Ecology Letters 14, 690–701.

Mcart, S.H., Cook-Patton, S.C. & Thaler, J.S. (2012). Relationships between arthropod richness, evenness, and diversity are altered by complementarity among plant genotypes. Oecologia 168, 1013–1021.

Mimura, M., Yahara, T., Faith, D.P., Vázquez-Domínguez, E., Colautti, R.I., Araki, H., Javadi, F., Núñez-Farfán, J., Mori, A.S., Zhou, S., Ollingsworth, P.M., Neaves, L.E., Fukano, Y., Smith, G.F., Sato, Y.-I., ET AL. (2016). Understanding and monitoring the consequences of human impacts on intraspecific variation. Evolutionary Applications 10, 121–139.

Morrissey, M.B. (2016a). Meta-analysis of magnitudes, differences and variation in evolutionary parameters. Journal of Evolutionary Biology 29, 1882–1904.

Morrissey, M.B. (2016b). Rejoinder: Further considerations for meta-analysis of transformed quantities such as absolute values. Journal of Evolutionary Biology 29, 1922–1931.

Naeem, S., Thompson, Lindsey J, Lawler, Sharon P, Lawton, John H & Woodfin, Richard M (1994). Declining biodiversity can alter the performance of ecosystems. Nature 368, 734–737.

Nakagawa, S. & Cuthill, I.C. (2007). Effect size, confidence interval and statistical significance: a practical guide for biologists. Biological Reviews 82, 591–605.

Nakagawa, S. & Santos, E.S.A. (2012). Methodological issues and advances in biological meta-analysis. Evolutionary Ecology 26, 1253–1274.

Nell, C.S., Meza-Lopez, M.M., Croy, J.R., Nelson, A.S., Moreira, X., Pratt, J.D. & Mooney, K.A. (2018). Relative effects of genetic variation sensu lato and sexual dimorphism on plant traits and associated arthropod communities. Oecologia 187, 389–400.

Noble, D.W.A., Lagisz, M., O’Dea, R.E. & Nakagawa, S. (2017). Nonindependence and sensitivity analyses in ecological and evolutionary meta-analyses. Molecular Ecology 26, 2410–2425.

Odling-Smee, J., Laland, K.N. & Feldman, M.W. (2003). Niche construction the neglected process in evolution. Princeton University Press.

Paine, R.T. (1969). A note on trophic complexity and community stability. The American Naturalist 103, 91–93.

Palkovacs, E.P., Fryxell, D.C., Turley, N.E. & Post, D.M. (2015). Ecological effects of intraspecific consumer biodiversity for aquatic communities and ecosystems. In Aquatic Functional Biodiversity pp. 37–51. Elsevier.

Palkovacs, E.P., Marshall, M.C., Lamphere, B.A., Lynch, B.R., Weese, D.J., Fraser, D.F., Reznick, D.N., Pringle, C.M. & Kinnison, M.T. (2009). Experimental evaluation of evolution and coevolution as agents of ecosystem change in Trinidadian streams. Philosophical Transactions of the Royal Society B-Biological Sciences 364, 1617–1628.

Palkovacs, E.P. & Post, D.M. (2009). Experimental evidence that phenotypic divergence in predators drives community divergence in prey. Ecology 90, 300–305.

Pinheiro, J., Bates, D., Debroy, S., Sarkar, D. & R CORE TEAM (2014). nlme: linear and nonlinear mixed effects models. R package version 3.1-117.

Post, D.M., Palkovacs, E.P., Schielke, E.G. & Dodson, S.I. (2008). Intraspecific variation in a predator affects community structure and cascading trophic interactions. Ecology 89, 2019–2032.

Pratt, J.D., Datu, A., Tran, T., Sheng, D.C. & Mooney, K.A. (2017). Genetically based latitudinal clines in *Artemisia californica* drive parallel clines in arthropod communities. Ecology 98, 79–91.

Pregitzer, C.C., Bailey, J.K. & Schweitzer, J.A. (2013). Genetic by environment interactions affect plant-soil linkages. Ecology and Evolution 3, 2322–2333.

Reiss, J., Bridle, J.R., Montoya, J.M. & Woodward, G. (2009). Emerging horizons in biodiversity and ecosystem functioning research. Trends in Ecology & Evolution 24, 505–514.

Renneville, C., Rouzic, A.L., Baylac, M., Millot, A., Loisel, S. & Edeline, E. (2016). Morphological drivers of trophic cascades. Oikos 125, 1193–1202.

Robinson, K.M., Ingvarsson, P.K., Jansson, S. & Albrectsen, B.R. (2012). Genetic variation in functional traits influences arthropod community composition in aspen (*Populus tremula L*.). PLoS ONE 7, e37679.

Rodriguez-Cabal, M.A., Barrios-Garcia, M.N., Rudman, S.M., Mckown, A.D., Sato, T. & Crutsinger, G.M. (2017). It is about time: genetic variation in the timing of leaf-litter inputs influences aquatic ecosystems. Freshwater Biology 62, 356–365.

RoyautÉ, R. & Pruitt, J.N. (2015). Varying predator personalities generates contrasting prey communities in an agroecosystem. Ecology 96, 2902–2911.

Rudgers, J.A. & Whitney, K.D. (2006). Interactions between insect herbivores and a plant architectural dimorphism. Journal of Ecology 94, 1249–1260.

Rudman, S.M., Rodriguez-Cabal, M.A., Stier, A., Sato, T., Heavyside, J., El-Sabaawi, R.W. & Crutsinger, G.M. (2015). Adaptive genetic variation mediates bottom-up and top-down control in an aquatic ecosystem. Proceedings of the Royal Society B: Biological Sciences 282, 125–132.

Rudman, S.M. & Schluter, D. (2016). Ecological impacts of reverse speciation in threespine stickleback. Current Biology 26, 490–495.

Rudolf, V.H.W. & Rasmussen, N.L. (2013a). Ontogenetic functional diversity: Size structure of a keystone predator drives functioning of a complex ecosystem. Ecology 94, 1046–1056.

Rudolf, V.H.W. & Rasmussen, N.L. (2013b). Population structure determines functional differences among species and ecosystem processes. Nature Communications 4.

Schmitz, O.J., Hawlena, D. & Trussell, G.C. (2010). Predator control of ecosystem nutrient dynamics. Ecology Letters 13, 1199–1209.

Schmitz, O.J., Raymond, P.A., Estes, J.A., Kurz, W.A., Holtgrieve, G.W., Ritchie, M.E., Schindler, D.E., Spivak, A.C., Wilson, R.W., Bradford, M.A., Christensen, V., Deegan, L., Smetacek, V., Vanni, M.J. & Wilmers, C.C. (2014). Animating the carbon cycle. Ecosystems 17, 344–359.

Schweitzer, J.A., Bailey, J.K., Fischer, D.G., Leroy, C.J., Lonsdorf, E.V., Whitham, T.G. & Hart, S.C. (2008). Plant–soil–microorganism interactions: heritable relationship between plant genotype and associated soil microorganisms. Ecology 89, 773–781.

Semchenko, M., Saar, S. & Lepik, A. (2017). Intraspecific genetic diversity modulates plant-soil feedback and nutrient cycling. New Phytologist 216, 90–98.

Senior, A.M., Grueber, C.E., Kamiya, T., Lagisz, M., O’Dwyer, K., Santos, E.S.A. & Nakagawa, S. (2016). Heterogeneity in ecological and evolutionary meta-analyses: its magnitude and implications. Ecology 97, 3293–3299.

Shipley, B. (2000). Cause and correlation in biology: a user’s guide to path analysis, structural equation and causal inference. Cambridge University Press.

Siefert, A., Violle, C., Chalmandrier, L., Albert, C.H., Taudiere, A., Fajardo, A., Aarssen, L.W., Baraloto, C., Carlucci, M.B., Cianciaruso, M.V., De L. Dantas, V., De Bello, F., Duarte, L.D.S., Fonseca, C.R., Freschet, G.T., ET AL. (2015). A global meta-analysis of the relative extent of intraspecific trait variation in plant communities. Ecology Letters 18, 1406–1419.

Silfver, T., Kontro, M., Paaso, U., Karvinen, H., Keski-Saari, S., KeinÄNen, M., Rousi, M. & Mikola, J. (2018). Intrapopulation genotypic variation in leaf litter chemistry does not control microbial abundance and litter mass loss in silver birch, *Betula pendula*. Plant and Soil 426, 253–266.

Silfver, T., Mikola, J., Rousi, M., Roininen, H. & Oksanen, E. (2007). Leaf litter decomposition differs among genotypes in a local *Betula pendula* population. Oecologia 152, 707–714.

Silfver, T., Paaso, U., Rasehorn, M., Rousi, M. & Mikola, J. (2015). Genotype x herbivore effect on leaf litter decomposition in *Betula Pendula* saplings: ecological and evolutionary consequences and the role of secondary metabolites. PLoS One 10.

Silfver, T., Rousi, M., Oksanen, E. & Roininen, H. (2014). Genetic and environmental determinants of insect herbivore community structure in a *Betula pendula* population. F1000Research 3, 34–34.

Southwood, T.R., Brown, V.K. & Reader, P.M. (1979). The relationships of plant and insect diversities in succession. Biological Journal of the Linnean Society 12, 327–348.

Sthultz, C.M., Whitham, T.G., Kennedy, K., Deckert, R. & Gehring, C.A. (2009). Genetically based susceptibility to herbivory influences the ectomycorrhizal fungal communities of a foundation tree species. New Phytologist 184, 657–667.

Tack, A.J.M., Ovaskainen, O., Pulkkinen, P. & Roslin, T. (2010). Spatial location dominates over host plant genotype in structuring an herbivore community. Ecology 91, 2660–2672.

Tack, A.J.M. & Roslin, T. (2011). The relative importance of host-plant genetic diversity in structuring the associated herbivore community. Ecology 92, 1594–1604.

Tovar-Sanchez, E., Valencia-Cuevas, L., Castillo-Mendoza, E., Mussali-Galante, P., Perez-Ruiz, R.V. & Mendoza, A. (2013). Association between individual genetic diversity of two oak host species and canopy arthropod community structure. European Journal of Forest Research 132, 165–179.

Trap, J., Hättenschwiler, S., Gattin, I. & Aubert, M. (2013). Forest ageing: An unexpected driver of beech leaf litter quality variability in European forests with strong consequences on soil processes. Forest Ecology and Management 302, 338–345.

Utsumi, S. (2015). Feeding evolution of a herbivore influences an arthropod community through plants: implications for plant-mediated eco-evolutionary feedback loop. Journal of Ecology 103, 829–839.

Vanni, M.J. (2002). Nutrient cycling by animals in freshwater ecosystems. Annual Review of Ecology and Systematics 33, 341–370.

Vellend, M. (2005). Species diversity and genetic diversity: parallel processes and correlated patterns. The American Naturalist 166, 199–215.

Vellend, M. & Geber, M.A. (2005). Connections between species diversity and genetic diversity. Ecology Letters 8, 767–781.

Violle, C., Enquist, B.J., Mcgill, B.J., Jiang, L., Albert, C.H., Hulshof, C., Jung, V. & Messier, J. (2012). The return of the variance: intraspecific variability in community ecology. Trends in Ecology & Evolution 27, 244–252.

Violle, C., Navas, M.-L., Vile, D., Kazakou, E., Fortunel, C., Hummel, I. & Garnier, E. (2007). Let the concept of trait be functional! Oikos 116, 882–892.

Wagg, C., Boller, B., Schneider, S., Widmer, F. & Van Der Heijden, M.G.A. (2015). Intraspecific and intergenerational differences in plant-soil feedbacks. Oikos 124, 994–1004.

Walsh, M.R., Delong, J.P., Hanley, T.C. & Post, D.M. (2012). A cascade of evolutionary change alters consumer-resource dynamics and ecosystem function. Proceedings of the Royal Society B-Biological Sciences 279, 3184–3192.

Wang, X.-Y., Miao, Y., Yu, S., Chen, X.-Y. & Schmid, B. (2014). Genotypic diversity of an invasive plant species promotes litter decomposition and associated processes. Oecologia 174, 993–1005.

Weis, J.J. & Post, D.M. (2013). Intraspecific variation in a predator drives cascading variation in primary producer community composition. Oikos 122, 1343–1349.

Werner, E.E. & Peacor, S.D. (2003). A review of trait-mediated indirect interactions in ecological communities. Ecology 84, 1083–1100.

Whitham, T.G., Bailey, J.K., Schweitzer, J.A., Shuster, S.M., Bangert, R.K., Leroy, C.J., Lonsdorf, E.V., Allan, G.J., Difazio, S.P., Potts, B.M., Fischer, D.G., Gehring, C.A., Lindroth, R.L., Marks, J.C., Hart, S.C., ET AL. (2006). A framework for community and ecosystem genetics: from genes to ecosystems. Nature Reviews Genetics 7, 510–523.

Whitham, T.G., Young, W.P., Martinsen, G.D., Gehring, C.A., Schweitzer, J.A., Shuster, S.M., Wimp, G.M., Fischer, D.G., Bailey, J.K., Lindroth, R.L., Woolbright, S. & Kuske, C.R. (2003). Community and ecosystem genetics: A consequence of the extended phenotype. Ecology 84, 559–573.

Wilkinson, A., Alexander, I. & Johnson, D. (2012). Genotype identity determines productivity and CO2 efflux across a genotype-species gradient of ectomycorrhizal fungi. Fungal Ecology 5, 571–580.

Wilkinson, A., Solan, M., Taylor, A.F.S., Alexander, I.J. & Johnson, D. (2010). Intraspecific diversity regulates fungal productivity and respiration. Plos One 5, e12604.

Zytynska, S.E., Fay, M.F., Penney, D. & Preziosi, R.F. (2011). Genetic variation in a tropical tree species influences the associated epiphytic plant and invertebrate communities in a complex forest ecosystem. Philosophical Transactions of the Royal Society B: Biological Sciences 366, 1329–1336.

Zytynska, S.E., Khudr, M.S., Harris, E. & Preziosi, R.F. (2012). Genetic effects of tank-forming bromeliads on the associated invertebrate community in a tropical forest ecosystem. Oecologia 170, 467–475.

